# WGCNA and Machine Learning for Screening Potential Biomarkers in traumatic brain injury

**DOI:** 10.1101/2023.08.29.555247

**Authors:** Yu Liu, Zongren Zhao, Jinyu Zheng

**Author notes:** Correspondence: Jinyu Zheng. Author Contributions: Yu Liu and Zongren Zhao contributed equally to this work.

## Abstract

**Background:** Traumatic brain injury (TBI) is more common than ever and is becoming a global public health issue. However, there are no sensitive diagnostic or prognostic biomarkers to identify TBI, which leads to long-term consequences. In this study, we aim to identify genes that contribute to brain injury and to identify potential mechanisms for its progression in the early stages.

**Method:** From the Gene Expression Omnibus (GEO) database, we downloaded GSE2871’s gene expression profiles. Weighted gene coexpression network analyses (WGCNA) were conducted on differentially expressed genes (DEGs), and the DEGs were analyzed by Gene Set Enrichment Analysis(GSEA). An enrichment analysis of Gene Ontology (GO) and Kyoto Encyclopedia of Genes and Genomes (KEGG) was performed for understanding the biological functions of genes. The potential biomarkers were identified using 3 kinds of machine learning algorithms. Nomogram was constructed using the “rms” package. And the receiver operating characteristic curve (ROC) was plotted to detect and validate our prediction model sensitivity and specifificity.

**Results:** Between samples with and without brain injury, 107 DEGs were identified, including 47 upregulated genes and 60 downregulated genes. On the basis of WGCNA and DEGs, 97 target genes were identified. In addition, biological function analysis indicated that target genes were primarily involved in the interaction of neuroactive ligands with receptors, taste transduction, cortisol synthesis and secretion, potassium ion transport. Based on machine learning algorithms, LOC103691092, Npw could be potentially useful biomarkers for TBI and showed good diagnostic values. Finally, a nomogram was constructed of the expression levels of these seven target genes to predict level of TBI, and the ROC showed that these genes can be used as hub genes after TBI.

**Conclusion:** LOC103691092, NPW, STK39, KCND3, APOC3, FOXE3, and CHRNB1 were identified as hub genes of TBI. These findings can provide a new direction for the diagnosis and treatment of TBI.

## Introduction

Traumatic brain injury (TBI) is a major global health problem and a leading cause of death and disability^1^. It occurs as a result of direct impact or impact to the head from factors such as motor vehicles, crush and assault^2^. Even non-fatal injuries can lead to severe lifelong disability, which has significant implications for the injured and their families, as well as for medical costs^3-5^. It is well known that the earlier a non-fatal traumatic brain injury is detected and treated, the less long-term impact it has on the injured person^6,7^. The severity of TBI was graded using the Glasgow Coma Scale (GCS), which was divided into mild, moderate and severe according to the degree of injury^8^. Mild traumatic brain injury (mTBI) has been reported to account for 70% to 90% of all traumatic brain injury cases^9^. In the absence of a diagnosis of mTBI, its effects can lead to cognitive impairment, depression, and headache, among others^10^. At present, clinical imaging and electroencephalogram detection methods are not obvious for the detection of some mTBI, which makes it easy to be confused with concussion, and thus fail to treat patients in time and correctly^11,12^. Distinct patterns of gene expression following traumatic brain injury will occur in a time- and injury-dependent fashion. In particular, changes in expression of enzymes involved in energy metabolism and neuroplasticity will be detected.

In this study, we performed gene expression level analysis on the downloaded dataset to obtain differentially expressed genes between TBI patients and normal subjects. Combined the downloaded data with WGCNA and machine learning algorithm, a total of 7 core genes were screened out, and these 7 core genes were used to construct the nomogram. Traumatic brain injury (TBI) induces a complex cascade of molecular and physiological effects. This study proposes to investigate the gene expression profile in cortex and hippocampus over early time points, following two different injury severities. These results will complement prior knowledge of both metabolic and neuroplastic changes after TBI, as well as serve as a starting point to investigate additional gene families whose expression is altered after TBI. Based on this, we can relatively accurately distinguish between the clinically difficult concussion and mTBI, and can also target these genes for treatment.

## Materials and methods Data

### Collection

The gene expression profile (GSE2871) was obtained from the GEO database (https://www.ncbi.nlm.nih.gov/geo/), which was sequenced using the GPL85 platform. At early post-injury timepoint, animals will be sacrificed, brain regions (parietal cortex and hippocampus, ipsilateral and contralateral to injury) will be dissected and RNA isolated(Supplementary figure 1). RNA will be used to synthesize cRNA probes for microarray hybridization. We created a Github page and uploaded the raw data and code(https://github.com/guanr80/guanr80.git).

### Identifification of DEGs and GSEA

The “Limma” R package was used to screen DE Gs between TBI and normal samples, and genes with P < 0.05 and |log2FC| >1 were regarded as DEGs. GSEA-4.1.0 was used to input the expression data and phenotypic data, and the five most significantly up-regulated pathways and the most significantly down-regulated pathways were plotted, respectively.

### Screening of the Critical Genes

To find out the core genes that were altered after TBI, the downloaded dataset was used to construct a weighted gene co-expression network using the “WGCNA” R package. To obtain an accurate network, we performed a cluster analysis of the samples. And then, we calculated the Pearson correlation coefficient between each pair of genes to evaluate the expression similarity of genes and acquire a correlation matrix. We further used a soft threshold function to transform the correlation matrix into a weighted neighborhood matrix, and a soft join algorithm was used to select the optimal soft threshold to ensure that gene correlations fit the scale-free distribution to the greatest extent possible. Subsequently, the neighborhood matrix was transformed into a topological overlap matrix(TOM). After obtaining the co-expression modules, the key modules were screened out by correlation analysis, and the genes of the key modules were regarded as the key genes.Based on WGCNA screening, DEGs and key genes were intersected to obtain the target genes.

### Functional Enrichment Analysis

R packages “clusterProfifiler” and “enrichplot” were used to perform GO assays and KEGG assays of DEGs with a statistically significant difference of at least P< 0.05.

### Identification of TBI hub genes based on machine learning algorithms

Least absolute shrinkage and selection operator (LASSO) is a regression analysis method that performs both gene selection and classification^13^. First, the R package glmnet (Version4.1.2) was used to fit the logistic LASSO regression model. Next, the SVM-RFE algorithm was used to screen potential genes using the “e1071” R package. In addition, the random forest (RF) algorithm was also conducted to screen potential genes using the “randomForest” R package. Finally, the intersection of the genes obtained by LASSO, SVM-RFE and RF machine learning algorithms was taken by Veen graph as the hub genes of TBI.

### Construction of the nomogram of TBI

We used “rms” in the R package to construct the nomogram out of the hub genes obtained by the machine learning algorithm. Next, ROC analysis was performed to evaluate whether hub genes could differentiate TBI samples from normal samples using the “pROC” R package.

### Analysis of associated genes of core genes and their potential drugs

We searched for associated genes from a public database (https://genemania.org). GeneMANIA uses extensive genomic and proteomic data to find functionally similar genes. We founded the relationship between drug targets (https://drugcentral.org/) and the main targets identified by this work.

### Evaluation of correlations and expression levels of key genes

Associations between hub genes were identified using Pearson correlation analysis, and P<0.05 was considered statistically significant. Wilcoxon’s rank-sum test was used to analyze the expression levels of hub genes.

## Results

### Identifification of DEGs

To explore biomarkers that are altered after TBI, this study retrospectively analyzed data on gene expression from TBI and normal samples in GSE2871 by setting the cut-off value as P < 0.05 and |log2FC| >1. 107 DEGs were identified, including 47 up regulated genes and 60 down regulated genes(Figure 1A, B). GSEA analysis was performed for differential genes to understand the main enrichment pathways of differential genes, and the top five up regulated pathways and the top five down regulated pathways were displayed(Figure 1C, D).

**FIGURE 1.**
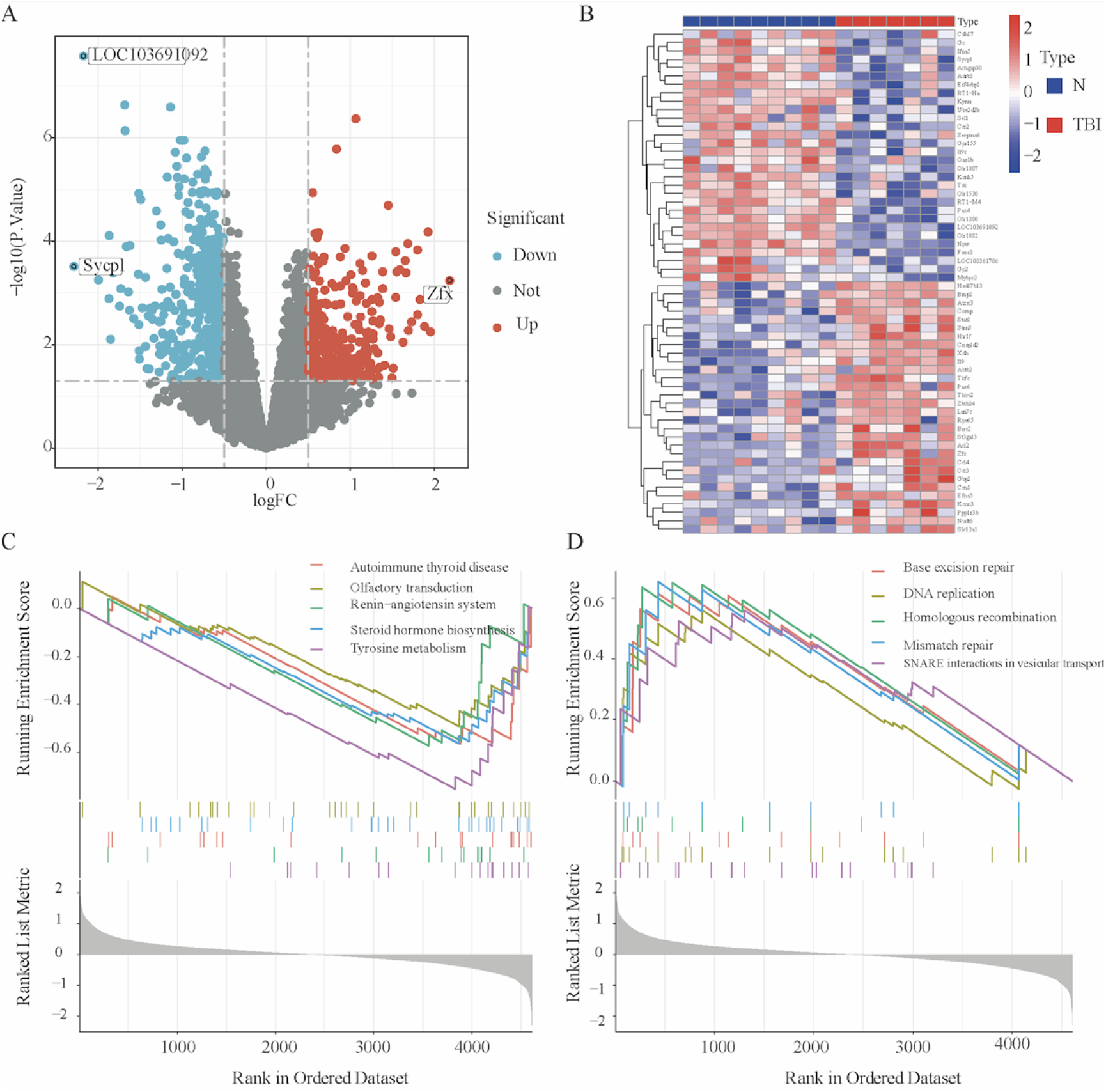
DEGs between TBI and normal samples. (A) Volcano plot of DEGs. (B) Heat map of DEGs. (C) GSEA of down-regulated DEGs. (D) GSEA of up-regulated DEGs.

### Screening of key modules and genes based on WGCNA

Analysis was performed to identify differentially expressed genes between TBI patients and normal controls. First of all, the soft threshold was selected for subsequent co-expression network construction (Figure 2A, B). The essence was to make the constructed network more consistent with the characteristics of scale-free networks. WGCNA was used to construct a co-expression network module and visually display the gene correlation of the modules. Co-expression modules were shown in a hierarchical cluster plot (Figure 2C). Multiple modules were shown to be associated with TBI through the moduletrait correlation studies. Each cell contains the corresponding correlation and P-value (Figure 2D). We show the association between module membership and gene importance using scatter plots (Figure 2E). The module “MEturquoise” had high association with TBI and was selected as TBI related module. By WGCNA screening, DEGs were crossed with key genes to obtain target genes (Figure 2F).

**FIGURE 2.**
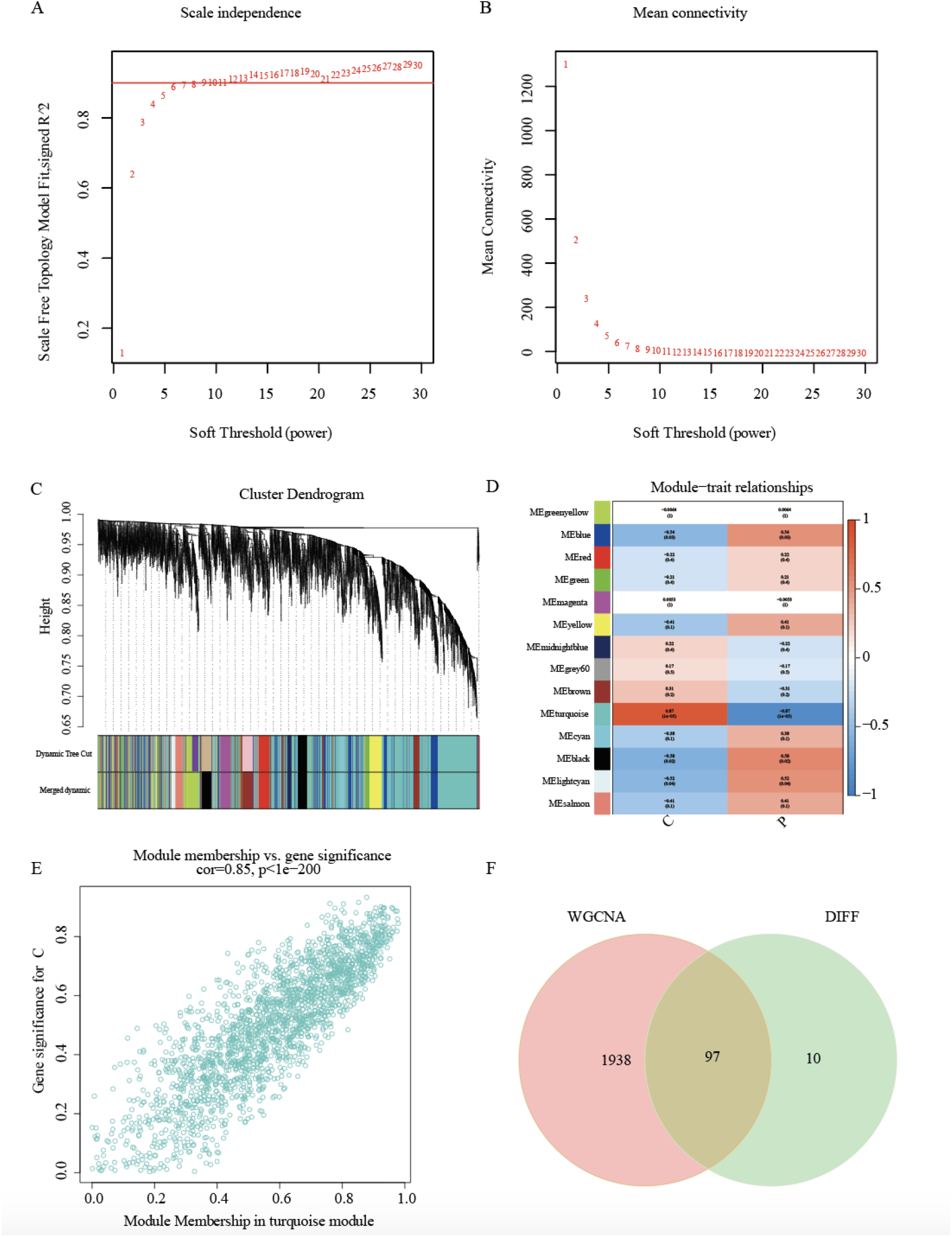
Construction of WGCNA modules. (A, B) The soft threshold was selected for subsequent co-expression network construction. (C) The cluster dendrogram of co-expression genes in TBI. (D) The module-trait relationship heat map. (E) Associations between module membership and gene importance is depicted in a scatter plot. (F) A Venn diagram was made to obtain the intersection of the target genes screened by the two methods.

### Functional Enrichment Analysis of DEGs

To further investigate the biological processes and signaling pathways associated with TBI DEGs, we utilized GO and KEGG analyses. The results of GO assays associated the most enriched biological process (BP) terms with potassium ion transport. The most enriched terms for cellular components (CC) were mainly associated with basolateral plasma membrane. The most enriched molecular function (MF) terms were associated with voltage-gated potassium channel activity (Figure 3A, B). The outcomes of KEGG assays revealed that DEGs were mainly enriched in pathways involved in neuroactive ligand-receptor interaction and cytokine-cytokine receptor interaction(Figure 3C, D).

**FIGURE 3.**
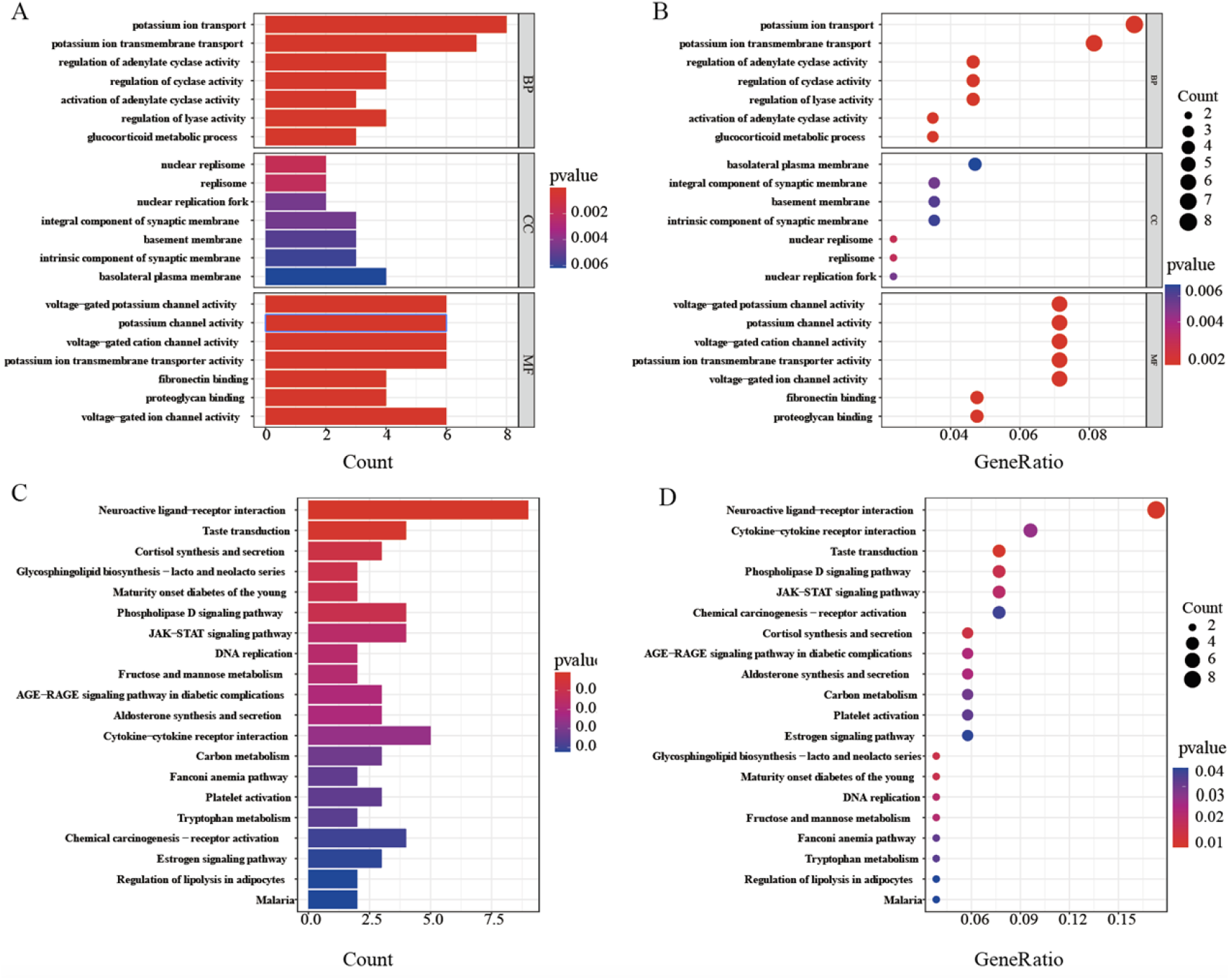
GO and KEGG analyses of target genes. (A, B) GO analyses of target genes. (C, D) KEGG analysis of target genes.

### Screening of key genes for TBI based on machine learning algorithms

To further identify the hub genes of TBI, we selected three machine learning algorithms to screen the target genes. The LASSO regression approach was used to narrow down the nine overlapping features, and nine variables were were further used in subsequent analyses(Figures 4A). The SVM-RFE analysis showed that a total of 86 potential genes were identified when the accuracy of SVM model was the best(Figures 4B). Meanwhile, the RF algorithm identified 32 genes at the lowest error rate(Figures 4C). Finally, seven hub genes changed after TBI were obtained according to the above three machine algorithms, which were STK39, KCND3, APOC3, FOXE3, CHRNB1, LOC103691092 and NPW(Figures 4D).

**FIGURE 4.**
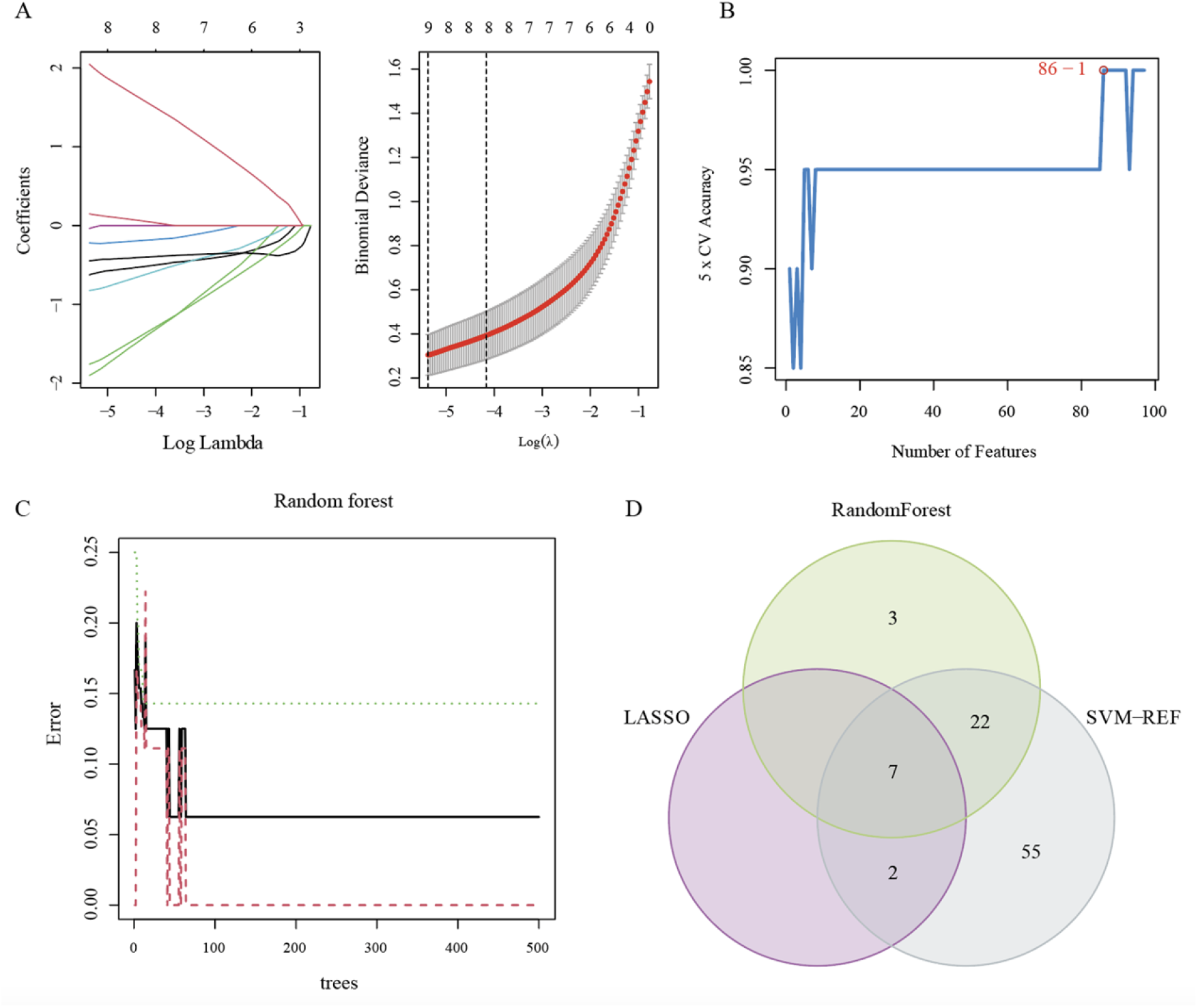
Identification of hub genes for TBI based on machine learning algorithms. (A) The Log (Lambda) value of the genes in the LASSO model and the most proper log (Lambda) value in the LASSO model. (B) The optimum accuracy rate of the SVM model based on the characteristic genes. (C) The RF module based on the characteristic genes. (D) The Venn diagram showing the overlapping genes in LASSO, SVM, and RF modules.

### Development of a nomogram for hub genes after TBI and diagnostic implications for the hub genes

Patients with “concussion” and “mild traumatic brain injury” are often encountered in clinical work, and the differentiation between the two is sometimes not so easy, and even some can not be distinguished on imaging^14^. At this time, some methods are needed to assist in diagnosis. A diagnostic nomogram was successfully constructed based on the eight genes for evaluating the incidence of TBI(Figure 5A). The nomogram based on the key genes derived by machine algorithms can solve the clinical difficult to distinguish between “concussion” and “mild traumatic brain injury”. To further investigate the role of hub genes in TBI, we first performed ROC analysis of these genes, which showed that these genes can be used as hub genes after TBI. These genes are LOC103691092, NPW, STK39, KCND3, APOC3, FOXE3 and CHRNB1(Figure 5B-H).

**FIGURE 5.**
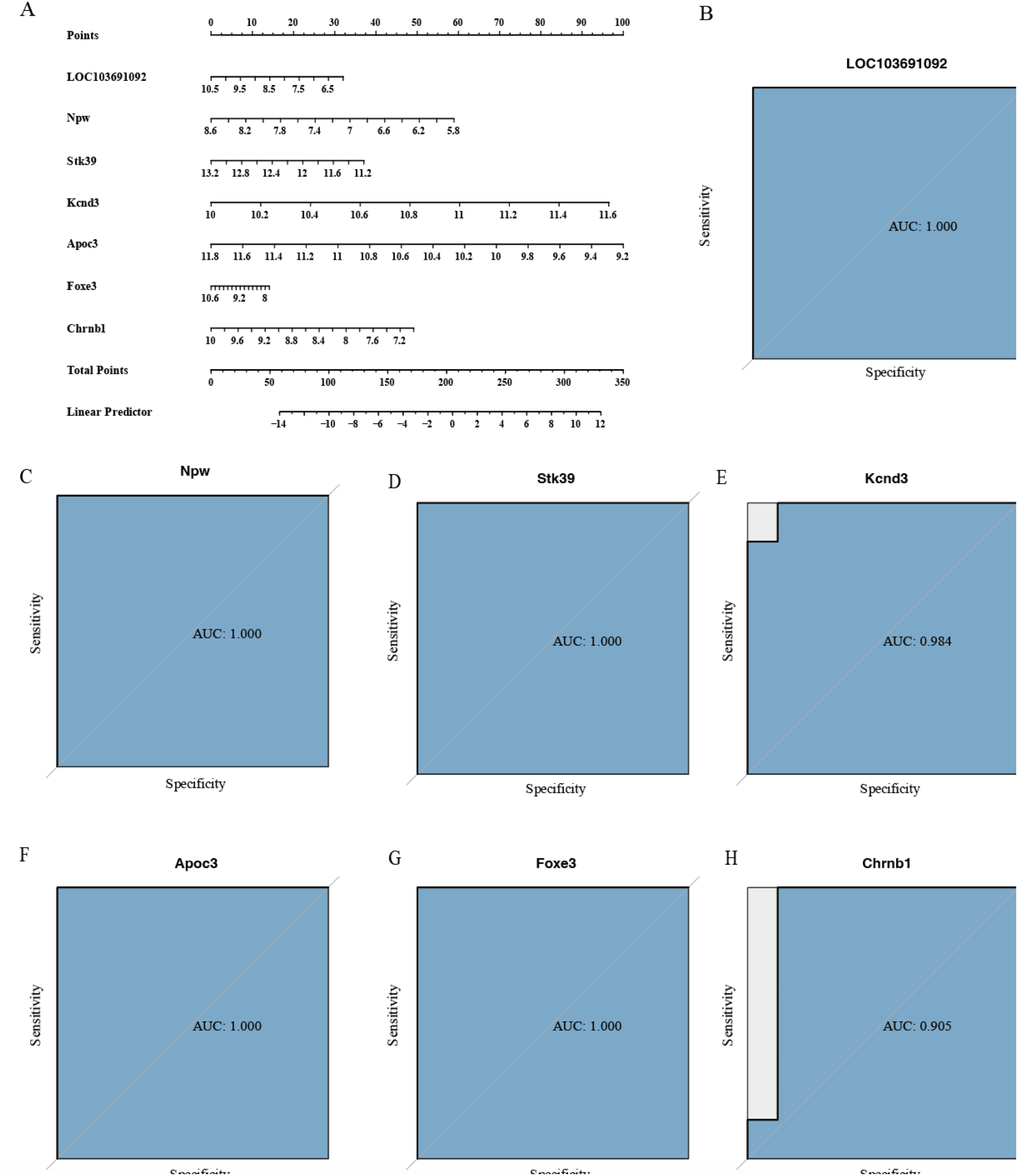
(A) A nomogram for hub genes after TBI. (B-H) ROC curves of hub genes of the dataset.

### Evaluation of correlations and expression levels of key genes

To further investigate the role of core genes in TBI, we observed their expression levels in TBI patients and normal samples. The results showed that among the seven hub genes, only KCND3 showed increased expression after TBI, while the others showed decreased expression(Figure 6A-G). After the differential expression levels of the 7 key genes were identified, correlation analysis was performed to further understand their relationship with each other. LOC103691092 was associated with the most genes, including STK39, KCND3 and CHRNB1(Figure 6H).

**FIGURE 6.**
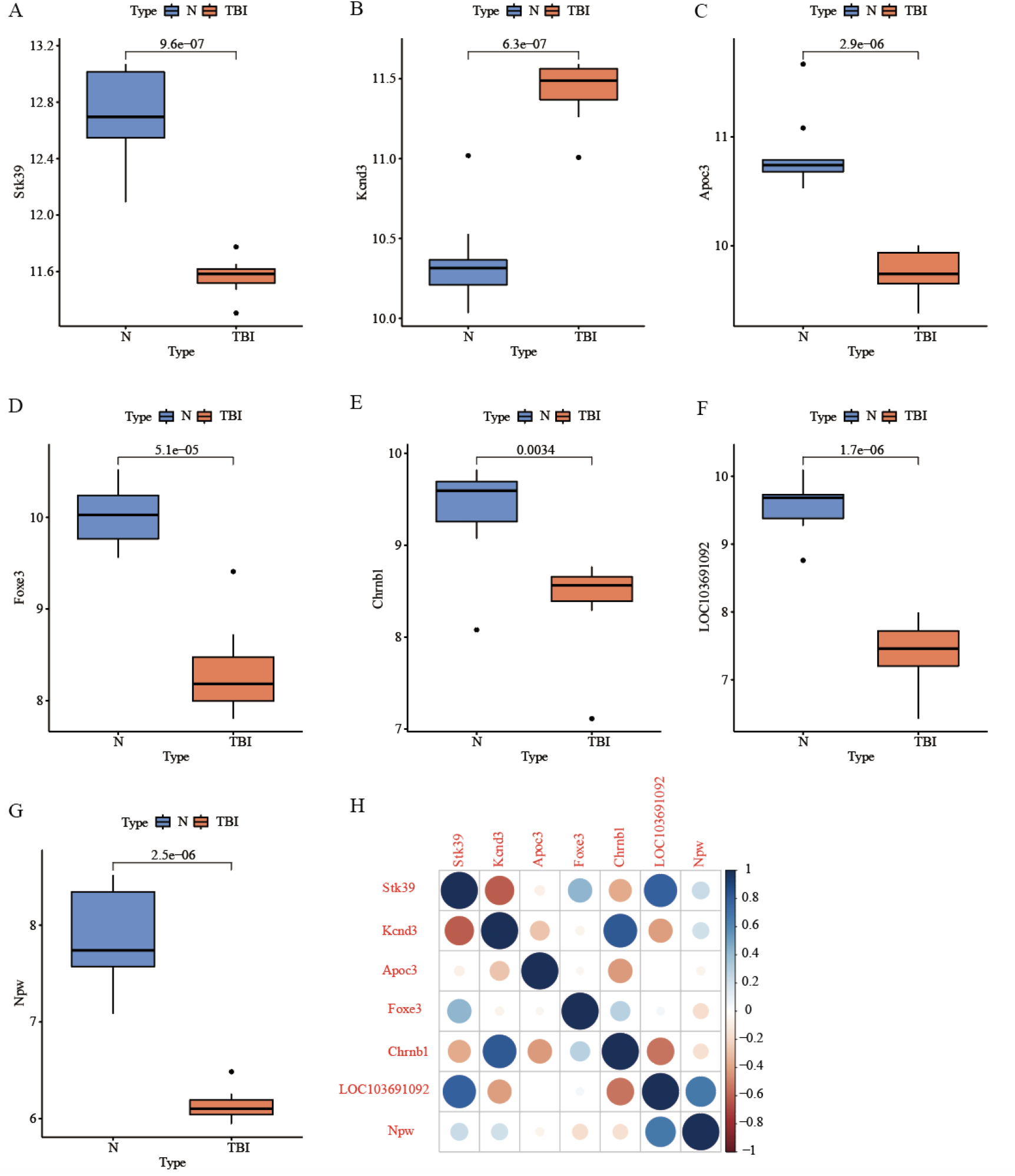
Evaluation of expression levels of key genes and their correlations. (A-G) The expression levels of hub genes in TBI patients and normal samples of the dataset. (H) Correlations of key genes.

### ssGSEA major pathways of core genes and their differential analysis in normal individuals and TBI

After the core genes that changed after TBI were analyzed, we performed ssGSEA analysis on them to display their main enriched pathways(Figure 7A). We then compared normal samples with major enriched pathways and TBI population for in-depth analysis of differential pathways(Figure 7B). Obviously, CHRNB1, LOC103691092 and STK39 are all closely related to “HALLMARK_PANCREAS_BETA_CELLS”. In addition to this, NPW has rich biological functions and is closely related to nine pathways. Among them, “HALLMARK_OXIDATIVE_PHOSPHORYLATION”, “HALLMARK_MYC_TARGETS_V1” and “HALLMARK_PROTEIN_SECRETION” had higher differential scores.

**FIGURE 7.**
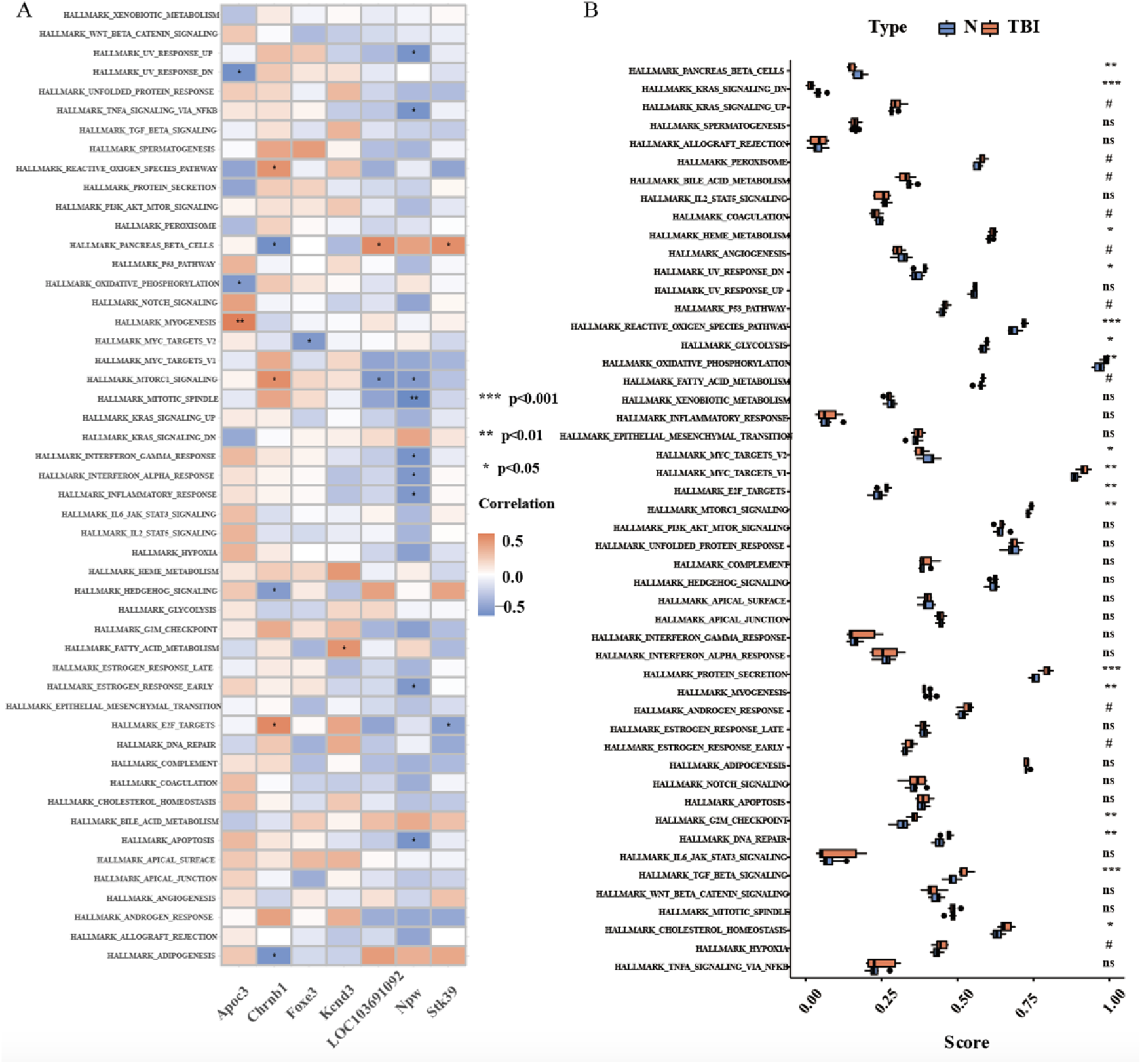
Major ssGSEA pathways of core genes and their differential analysis. (A) ssGSEA of the seven core genes. (B) Differential expression of core gene-enriched pathways.

### Analysis of associated genes of core genes and their potential drugs

After identifying seven TBI-related genes in this study, we identified the genes closely related to them from the GeneMANIA database(Figure 8A). The pathophysiological changes after TBI included ROS production, edema, inflammation, angiogenesis and metabolic related changes. We performed ssGSEA on the first seven genes, and the results showed that these genes were closely related to the pathophysiology of TBI(Figure 8B). The formation of ROS is closely related to FOXE3, CHENB1 and APOC3. Otherwise, APOC3 is closely related to inflammation caused by TBI. We identified a number of potential drugs that may act on some of the TBI-associated genes identified in this study in the public database (https://drugcentral.org/).

**FIGURE 8.**
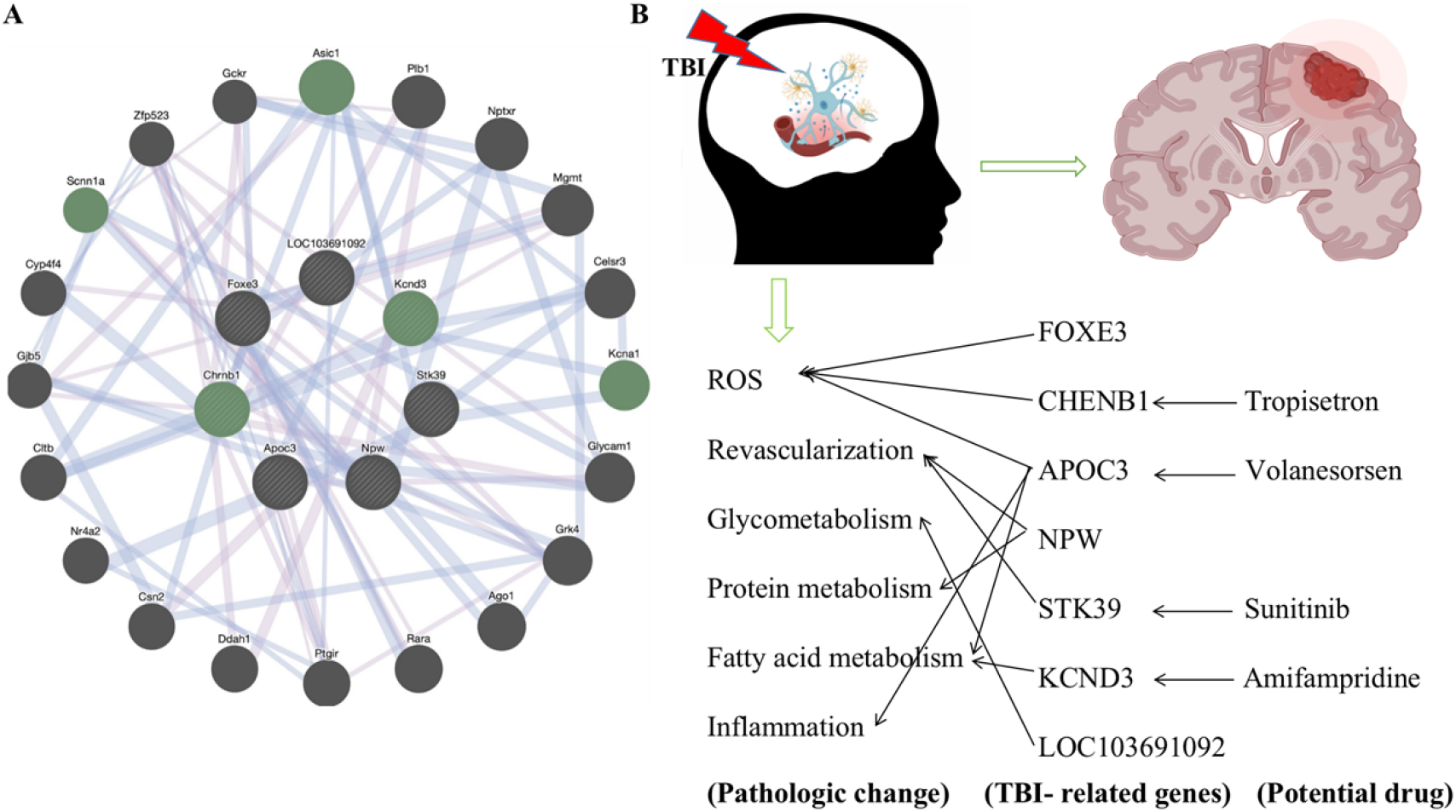
Analysis of associated genes and potential drugs. (A) The associated genes of key genes identified in this study. (B) Differential expression of core gene-enriched pathways.

## Discussion

TBI is a global public health problem that not only affects the long-term cognitive, physical, and mental health of patients, but also has a significant impact on families and caregivers^15^. Due to the lack of early imaging features in some mTBI patients, although mTBI patients have a history of brain trauma, it is often difficult to distinguish them from concussion, which often leads to delayed treatment and affects early treatment^16-19^. Therefore, biomarkers are needed to differentiate concussion from mTBI at an early stage in order to improve outcomes.

In the present study, we first obtained DEGs of TBI, and WGCNA screened key genes for the key module. Ninety-seven target genes were screened out by intersecting DEGs with key genes. Interestingly, these 97 target genes were mostly related to the interaction of neuroactive ligands with receptors, taste transduction, cortisol synthesis and secretion and potassium ion transport. Thus, we speculated that these genes might play key roles in TBI by regulating the interaction of neuroactive ligands with receptors, taste transduction, cortisol synthesis and secretion and potassium ion transport. Therefore, our study may contribute to understanding the molecular mechanisms underlying TBI.

Also, we identified LOC103691092, NPW, STK39, KCND3, APOC3, FOXE3 and CHRNB1 as hub genes using LASSO logistic regression, SVM-RFE and RF algorithms. Members of the NPW signaling system have been primarily detected and mapped to the CNS, and this signaling system has a wide range of functions, including regulation of inflammatory pain and neuroendocrine functions^20, 21^. Therefore, NPW may play an important role in the inflammatory pain and neuroendocrine process of TBI. Studies have shown a significant association between STK39 polymorphism and hypertension susceptibility, which also suggests that STK39 may be closely related to hypertension after TBI^22, 23^. Clinically, TBI patients often have electrophysiological disorders, and KCND3(the potassium voltage-gated channel subfamily D member 3) is closely related to electrophysiological balance, so this gene may play an important role in the process of TBI^24, 25^. Studies have shown that APOC3(apolipoprotein C-3) is involved in the regulation of vascular endothelium^26, 27^. CHRNB1 has been reported to be associated with inotropic regulation^28, 29^. LOC103691092 is a newly discovered gene and has not been reported.

Finally, we performed ssGSEA analysis on the 7 core genes, analyzed the highly enriched pathways, and compared the differences between normal samples and TBI samples. CHRNB1, LOC103691092 and STK39 are all closely related to “HALLMARK_PANCREAS_BETA_CELLS”. A number of studies have shown that this pathway is closely related to the regulation of human blood glucose levels, which also reasonably explains the mechanism of blood glucose elevation after brain injury^30-32^. The pathways with higher enrichment scores were “HALLMARK_OXIDATIVE_PHOSPHORYLATION”, “HALLMARK_MYC_TARGETS_V1” and “HALLMARK_PROTEIN_SECRETION”. These pathways contain potential mechanisms for the treatment of TBI, which remain to be further studied.

In conclusion, we identified seven genes that are closely related to TBI. Therefore, our study may help to understand the mechanism of TBI and identify TBI in head trauma patients. In addition, these key genes may also contribute to the targeted therapy of TBI, targeting specific genes to treat severe TBI that does not respond well to conventional methods. Of course, the role of hub genes remains to be further verified.

## Authors’ Contributions

Yu Liu and Zongren Zhao, conducted an evaluation of the information. Yu Liu collaborated in the preparation and correction of the work. Yu Liu wrote the manuscript, Zongren Zhao typesetted the images, and Jinyu Zheng revised and reviewed the article as a whole.

